# Hierarchy-guided Neural Networks for Species Classification

**DOI:** 10.1101/2021.01.17.427006

**Authors:** Mohannad Elhamod, Kelly M. Diamond, A. Murat Maga, Yasin Bakis, Henry L. Bart, Paula Mabee, Wasila Dahdul, Jeremy Leipzig, Jane Greenberg, Brian Avants, Anuj Karpatne

## Abstract

1. Species classification is an important task that is the foundation of industrial, commercial, ecological, and scientific applications involving the study of species distributions, dynamics, and evolution.
2. While conventional approaches for this task use off-the-shelf machine learning (ML) methods such as existing Convolutional Neural Network (ConvNet) architectures, there is an opportunity to inform the ConvNet architecture using our knowledge of biological hierarchies among taxonomic classes.
3. In this work, we propose a new approach for species classification termed Hierarchy-Guided Neural Network (**HGNN**), which infuses hierarchical taxonomic information into the neural network’s training to guide the structure and relationships among the extracted features. We perform extensive experiments on an illustrative use-case of classifying fish species to demonstrate that **HGNN** outperforms conventional ConvNet models in terms of classification accuracy, especially under scarce training data conditions.
4. We also observe that **HGNN** shows better resilience to adversarial occlusions, when some of the most informative patch regions of the image are intentionally blocked and their effect on classification accuracy is studied.

## Introduction

Depicting the branching pattern of taxa, phylogeny represents a hypothesis of evolutionary relationships based on shared similarities derived from common ancestry (Hennig, 1966). From conservation to zoology, phylogenetic relationships are critical for interpreting study results and implications in the biological sciences. One area, however, where this hierarchical information has yet to be fully incorporated is that of machine learning and image classification. Deep neural networks have found immense success in image classification problems with state-of-the-art ConvNet models (e.g., GoogleNet (Szegedy et al., 2015), AlexNet (Krizhevsky et al., 2012), and VGGNet (Simonyan and Zisserman, 2014)) reaching unprecedented performance on large-scale benchmark datasets such as ImageNet (Deng et al., 2009) and CIFAR (Krizhevsky, 2009). By design, deep neural networks function similarly to phylogenetic analyses by extracting a hierarchy of simpler to more complex forms of abstraction in hidden layers—simpler features at lower depths (e.g., edges and texture) are non-linearly composed to form complex features at higher depths (e.g., eyes and fins). This has motivated several recent architectural innovations in deep learning such as ResNet (He et al., 2016), ResNeXt (Xie et al., 2017), and DenseNet (Huang et al., 2017), that have enabled the learning of deep and complex hierarchy of hidden features. However, the innate hierarchy extracted by neural networks from data is not necessarily tied to known evolutionary relationships in real-world applications. In this work, we explore the question: *Is it possible to make use of known phylogenetic classes to inform the learning of features, and can it lead to better generalization and robustness?*

Image classification in real-world biological problems such as species classification is fraught with several challenges that limit the usefulness of state-of-the-art deep learning methods trained on benchmark datasets. First, real-world images of specimens suffer from various data quality issues such as damaged specimens and occlusions of key morphological features (Fox and Hartman, 2019), which can crucially impact classification performance. Figure 1 shows some relevant examples. Second, real-world datasets for classification are limited in their scale in comparison to benchmark datasets, with limited representative power in terms of number of species (Rathi et al., 2018; Ogunlana et al., 2015; Costa et al., 2013; Larsen et al., 2009; Lee et al., 2008; Allken et al., 2019; Rauf et al., 2019; Ding et al., 2017), or number of images per species (Rodrigues et al., 2010; Lee et al., 2003). This is especially true for rare species (Villon et al., 2021). Third, the hierarchy of features extracted by conventional deep learning frameworks, while useful for prediction, do not conform to known biological hierarchies and hence do not directly translate to advancing scientific knowledge, which is often a more important goal than improving predictive performance for a scientist (Karpatne et al., 2017). While these challenges are applicable to species classification problems involving a variety of taxa, in this study we focus on the problem of classifying the species of a fish specimen given a 2D image. We selected fishes for our study because they are a highly diverse, well-studied, and an ancient group of animals that comprise almost half of all vertebrate species (Helfman et al., 2009). Further, the phylogenetic relationships of fishes are well-studied (Betancur-R et al., 2017; Hughes et al., 2018), and the taxonomic classification of fishes is generally aligned with phylogeny.

**Fig. 1:**
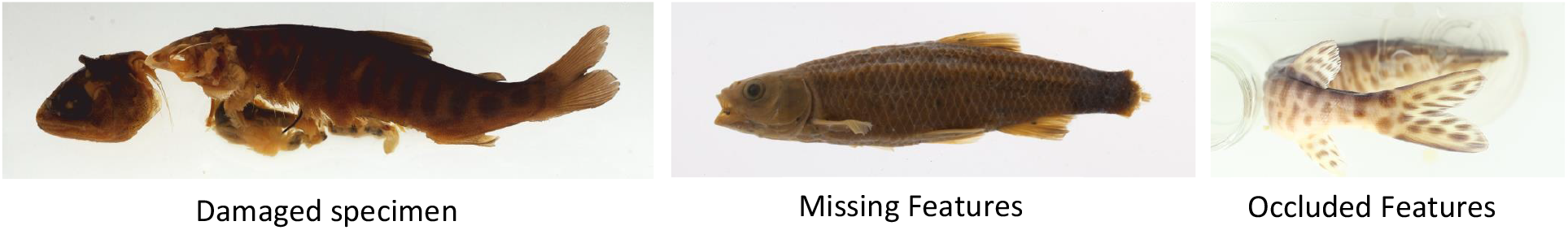
Fish images from museum collections, demonstrating the challenges of curating fish image datasets.

Early work on automated fish classification used basic computer vision and image processing techniques to extract shape features such as landmarks and measurements and used tools such as decision trees, discriminant function analysis, and support vector machines to classify species based on these features (Lee et al., 2003, 2008; Larsen et al., 2009; Ogunlana et al., 2015). Others have applied scale-invariant feature transform (SIFT) and principal component analysis (PCA), and then used nearest neighbor search for classification (Rodrigues et al., 2010). Only recently has the use of raw image features in its intrinsic high-dimensionality become more feasible, likely because of advances in computational capabilities. For example, (Hasija et al., 2017) employed graph-embedding discriminant analysis, which reduces the image set matching problem to a point-to-point classification problem.

Advances in computing power have also enabled researchers to use more flexible and powerful classification methods such as ConvNets, especially designed to work with high-dimensional images. The basic idea of a ConvNet is to learn convolutional kernels (or filters) of a fixed size at every layer, that are applied to the input image to generate multiple channels of image outputs for the next layer, followed by a final block of a max-pooling layer and a softmaxed fully connected layer to return class labels (Goodfellow et al., 2016). The number of feature maps is referred to as the width of the ConvNet, while the number of layers is termed as its depth. To further boost ConvNet’s performance, image preprocessing techniques can be used. For example, (Rathi et al., 2018) pre-processed the fish images by means of Gaussian blurring, erosion and dilation and Otsu thresholding (Otsu, 1979).

More recently, researchers have taken advantage of state-of-the-art architectures available in the field of deep learning for biological classification. For example, in a work by (Rauf et al., 2019), the technique of *transfer learning* was explored for fish classification, where neural network models *pre-trained* over large and diverse benchmark datasets were used as building blocks and then *fine-tuned* on the fish images. Transfer learning eliminates much of the arduous task of hyper-parameter tuning otherwise required in the field of deep learning, and allows researchers to build on top of well-tested benchmark neural network models. It also saves model development time and boosts classification performance, especially when the available task-specific training sets are small (Yosinski et al., 2014). This technique has already been successfully applied in other prior works on fish classification (Siddiqui et al., 2018; Allken et al., 2019) and fish detection (Salman et al., 2019).

Extensions of ConvNets have also been used for several tasks such as fish detection, counting, and classification. For example, (Salman et al., 2019) have used R-CNNs (Girshick et al., 2014) along with background subtraction and optical flow features to detect fish in underwater videos. Similarly, (Jalal et al., 2020) attack the problems of fish detection and classification using a YOLO deep neural network (Redmon et al., 2016) combined with a mixture of Gaussians model and optical flow features. In a different approach, (Villon et al., 2020) post-process the prediction of a deep learning model with confidence thresholding to obtain a misclassification risk estimation, which is particularly useful for identifying rare species. Finally, (Villon et al., 2021) have proposed using few-shot learning (Wang et al., 2020) to achieve better results on rare species. This, however, is at the expense of less robustness at distinguishing species that look too similar.

Our current method aims for a generic method that incorporates hierarchy to improve neural network models. Here we use taxonomic relationships from fish classification to serve as an example training dataset. Specifically, we present a novel deep learning architecture termed Hierarchy-Guided Neural Network (**HGNN**) that incorporates known hierarchy among classes (available as a two-level taxonomy: genus and species) to guide the learning of features at the hidden layers of the neural network. This work builds on a history of multi-label and hierarchical classification techniques using pre-built taxonomies (Silla and Freitas, 2011; Zhang and Zhou, 2013). Our proposed architecture shown in Figure 2 consists of two sub-modules (top and bottom rows) of ResNet models operating in parallel. We use the ResNet architecture in our work because it is currently among the most widely-used and best-performing ConvNet models for benchmark computer vision problems, including fish identification (Khan et al., 2020; Jalal et al., 2020; Villon et al., 2020; Ditria et al., 2020a), although our proposed idea of **HGNN** is generic and can work with any deep learning architecture. In Figure 2, the top row ResNet predicts the species class **s** of the input fish image **x**, while the bottom row predicts the genus class **g**. These ResNets learn a hierarchy of features (from simple to complex) at their hidden layers useful for the tasks of species and genus classification, respectively. While both these sub-modules can be viewed as learning separate features, we know that the genus features learned in the bottom ResNet represents features at a higher level of abstraction that are directly useful for the task of species classification. Building upon this knowledge in our proposed **HGNN** framework, we harness the genus features learned at an intermediate depth *H*_*g*_ of the genus sub-module, and aggregate them with the species features learned at the *H*_*s*_ depth of the species sub-modules. The combination of both species and genus features is then used for the task of species prediction.

**Fig. 2:**
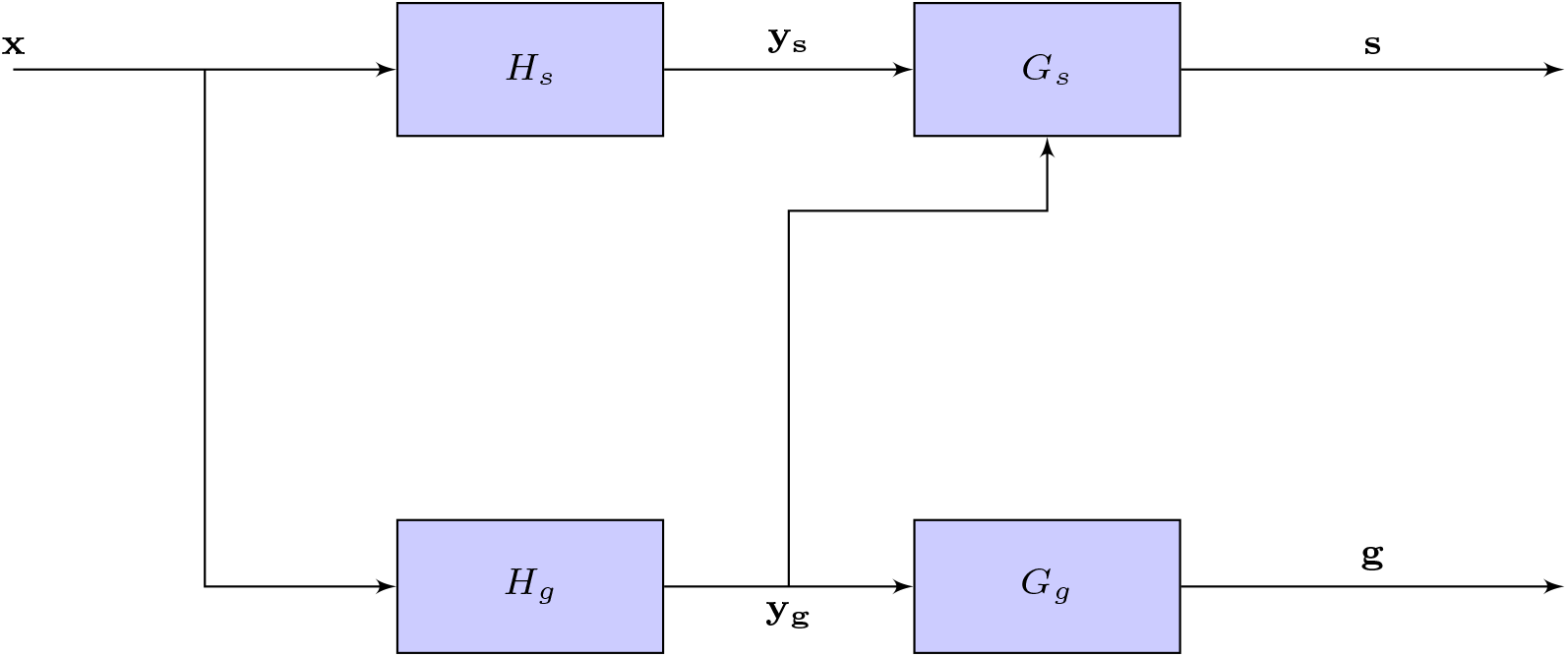
Schematic diagram of **HGNN**. The top ResNet predicts the species (**s**) of the input fish image (**x**), while the bottom ResNet predicts the genus (**g**). To leverage the relationship between genus and species classes for guiding the hidden features of our neural network, we harness the genus features learned at an intermediate depth (**y**_**g**_) of the genus ResNet and aggregate them with the species features learned at the **y**_**s**_ level of the species ResNet. The combination of both species and genus features are then used to make species class predictions. This architecture is described in details in the Materials and Methods section.

While using taxonomic information for automated fish classification is not novel (Kutlu et al., 2017), to our knowledge, the only body of work that has researched it before in the context of deep learning is by (dos Santos and Gonçalves, 2019). However, our proposed method is distinguished in two ways. First, while they have used the family and order information, we use the genus information. We argue that incorporating the genus yields more information gain as it involves more discriminative features than the order and family. Second, their model only uses the taxonomic information in the last fully-connected layer, while our philosophy is to use it at a convolutional level of the network as that allows for capturing localized visual features that are taxonomically plausible.

We demonstrate the effectiveness of our proposed **HGNN** model in learning meaningful, diverse, and robust features at the hidden layers of the neural network leading to better generalization performance in the target application of fish species classification, even in the paucity of training data. We also empirically test the robustness of our model to synthetically generated image occlusions, where salient regions of the input images were intentionally occluded to adversely affect classification performance. We observe that by anchoring our learned features to the biologically known hierarchy among genus and species classes, our model is much more robust to occlusions as compared to a data-only ‘black-box’ model that only uses image data and predicts the species with no genus information (i.e. using only the top ResNet in Figure 2).

## Materials and Methods

### HGNN framework

We first present our proposed **HGNN** architecture that incorporates hierarchy among genus and species classes in neural network construction. We consider the problem of predicting the target species **s** given input image **x** using a composition of neural network layers. We are also given the genus level class **g** for every input **x**.

We make two observations to motivate our proposed **HGNN** framework. First, we assume that the hierarchical taxonomy of genus and species classes captures a notion of derived similarity in terms of the discriminatory input features of every class. This is true, as illustrated in Figure 3, in the context of fish classification because species classes that belong to the same genus are more closely related phylogenetically than species classified in different genera. In the case of the species and genera analyzed here, with only a few exceptions, this is the case (Supporting Information, Table **??**). As a result, species that map to the same genus **g** should generally share similar features at the internal representation of the neural network (e.g., filters learned at the convolutional layers). This observations seems to align with some earlier work (dos Santos and Gonçalves, 2019). Second, while the mapping from **s** to **g** is one-to-one, the inverse mapping from **g** to **s** is not unique. Hence, along with the shared features learned for every **g**, we also need to learn unique features for every **s** to differentiate between species belonging to the same genus.

**Fig. 3:**
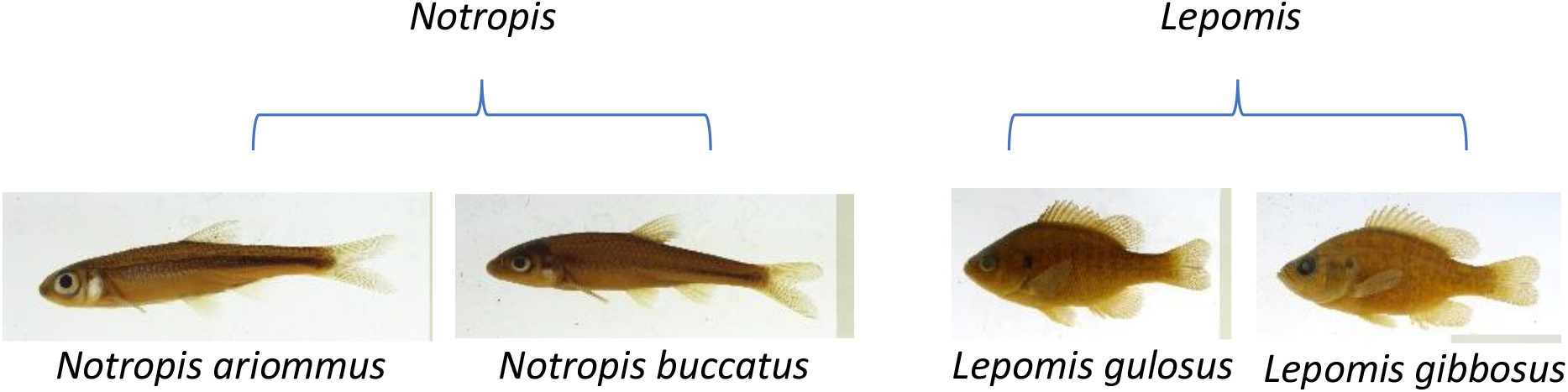
Species that belong to the same genus exhibit features that are similar because of common ancestry.

Building upon these two observations, we consider the following architectural composition of our neural network as shown in Figure 2. First, we use a functional block of layers *H*_*g*_ to extract hidden features at some intermediate depth of the neural network that are useful for predicting **g** as well as **s**. These hidden features are passed to another functional block *G*_*g*_ that predicts **g**. The complete chain of function compositions from **x** to **g** can be represented as *F*_*g*_(**x**), where *F*_*g*_ = *G*_*g*_ ○ *H*_*g*_ and ○ represents the function composition operator. Second, we learn another functional block *H*_*s*_ that extracts hidden features unique to every species. Finally, the features from *H*_*s*_ and *H*_*g*_ are combined using matrix addition and fed to another functional block of layers, *G*_*s*_ that predicts the target species **s**. The composition of functions mapping **x** to **s** can thus be given by *f* (**x**), where *F* = *G*_*s*_ ○ (*H*_*g*_ + *H*_*s*_).

To train the functional blocks in the complete **HGNN** architecture, we consider minimizing the following objective function:

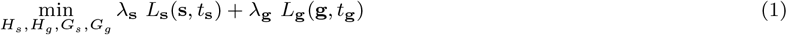

where *L*_**s**_ and *L*_**g**_ are loss (or error) functions defined on the space of species labels and genus labels, respectively, on the training set. Specifically, these loss functions act as a measure of difference between the correct classification (*t*_**s**_ and *t*_**g**_), and the prediction (**s** and **g**) on the training samples, respectively. We used the cross-entropy function as our preferred choice of loss function. Further, *λ*_**s**_ and *λ*_**g**_ are trade-off hyper-parameters balancing the relative importance of *L*_**s**_ and *L*_**g**_, respectively; their values are automatically assigned using the adaptive smoothing algorithm proposed in (Murugesan et al., 2016). Both the softmaxed outputs of our neural network model, **s** and **g**, are probability vectors whose entries range from 0 to 1 proportional to the model’s credence about each species and genus class, respectively.

As mentioned in the Introduction, our model is composed of two identical ResNets. The first ResNet comprises of *H*_*g*_ and *G*_*g*_, while *H*_*s*_ and *G*_*s*_ constitute the other. In our experiments, we found that the best point to extract the intermediate genus features (i.e. the point between *H*_*g*_ and *G*_*g*_) is right before the final max-pooling layer. The same point in the other ResNet is used to combine the genus and species features. Instead of initializing our neural network parameters (or weights) with arbitrary values, we used pre-trained weights of ResNet trained on the ImageNet benchmark dataset as a good starting solution for our target problem of fish classification. Then, by optimizing the loss function in equation (1) on the fish training dataset of interest, we fine-tuned the parameters of the entire network to be more specialized for our target task. This technique, which is called transfer learning (Tan et al., 2018a), is widely adopted in the field of deep learning particularly in applications of computer vision, and has proven its effectiveness in scenarios with data paucity. In our preliminary experiments, as shown in Figure 4, we have found using this mode of transfer learning to increase the model’s average performance by about 35%.

**Fig. 4:**
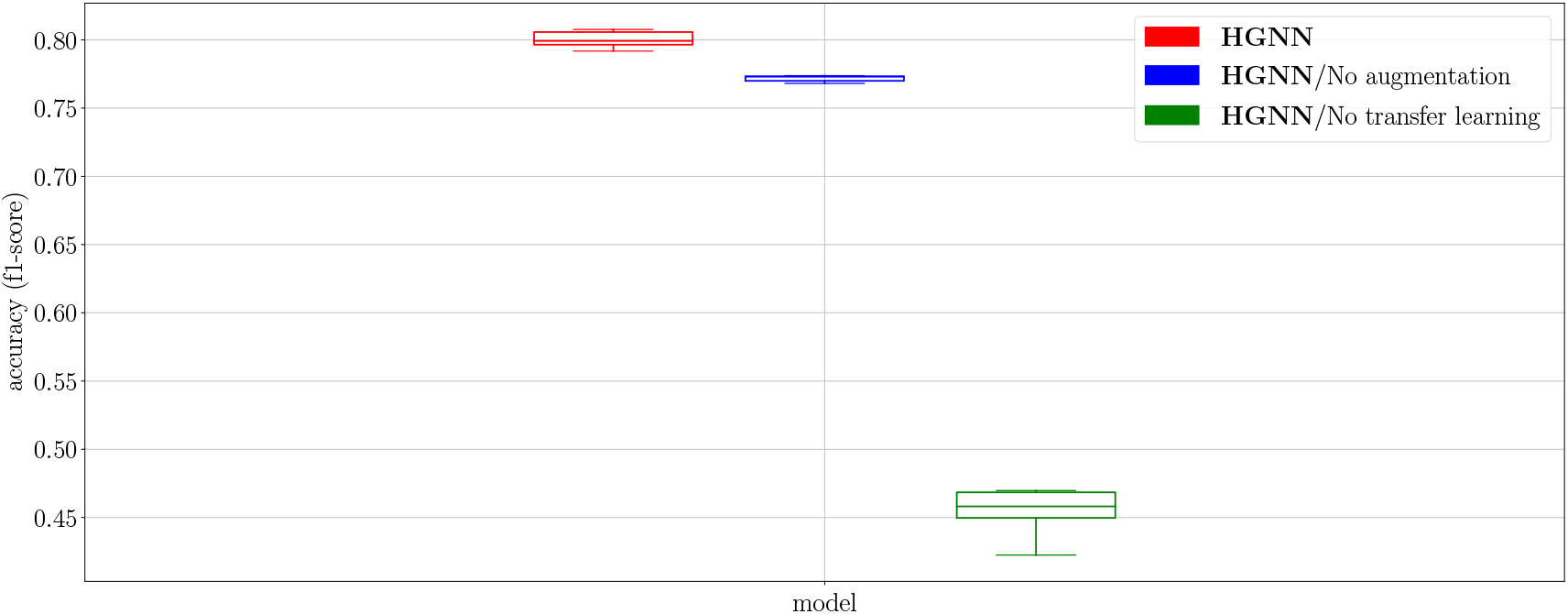
Comparison among different models, showing the impact of data augmentation and transfer learning on the classification performance of **HGNN** models.

## Evaluation

### Data Collection and Pre-processing

Our dataset comprises of images contributed by five museums that participated in the Great Lakes Invasives Network Project (GLIN). More information about this project can be found in the Data Availability section. This dataset, as is typical for biological species images, is highly imbalanced; some species have only a few images, while others have thousands. To alleviate this problem, and for computational feasibility, we created a number of subsets of the dataset for the purpose of training and evaluation. Specifically, we created two subsets that differ in terms of classification complexity (or difficulty). The first subset is called **Easy** and comes from a single museum (Illinois Natural History Survey). Therefore, its images are homogeneous in terms of lighting and camera conditions. The second is called **Hard** and its images are aggregated from across all museums, making it a larger, more diverse, and more complex dataset. Comparing results from these two datasets helps illustrate the effects of dataset complexity on classification performance. We further created two subsets of the **Easy** dataset by capping the number of images per species in the **Easy** dataset to 50 or 100. These different dataset sizes help illustrate how training data paucity impacts the model’s classification performance. Henceforth, the suffix of the datset will refer to the number of images per species. For example, **Easy**/100 has 100 images per species. Table 1 gives a statistical summary of each dataset considered in this study. More details can be found in the Supporting Information document, Tables **??, ??**, and **??**

**Table 1.** Statistics of the subsets of the GLIN dataset used in this study for training and evaluation.

| Dataset | # of images | # of species | # of genera | # of images per species |
| --- | --- | --- | --- | --- |
| GLIN (All) | 63758 | 575 | 187 | 1 to 7935 |
| **Hard** | 4882 | 102 | 26 | 30 to 50 |
| **Easy**/100 | 3762 | 38 | 11 | 63 to 100 |
| **Easy**/50 | 1900 | 38 | 11 | 50 |

The acquired fish images typically contained a ruler, specimen label(s), and species tags along with the fish specimen. To retain only the fish region in the images, we trained a 2D Unet model (Goodfellow et al., 2016) using a small portion of our data in the ANTsRNet software (Tustison et al., 2018). We manually segmented the background, fish, scale bar, and field notes on 550 images using 3D Slicer (Kikinis et al., 2014). We used weights from the trained model to automatically mask and crop the fish specimen portion of the remainder images. With the exception of rare cases where the fish overlapped the scale bar and/or the field notes, which were discarded, this pipeline resulted in successful generation of RGB fish-only images at the original resolution. The pipeline was implemented in R using ANTsR (Avants, 2019) and ANTsRNet.

Once the cropped fish images were obtained, we performed data augmentation by randomly applying standard image transformations used in deep learning for computer vision, including translations of up to 0.25 of the image dimension, flips with a probability of 30%, rotations of up to 60°, and Gaussian random intensity variations using PCA with *σ* = 0.1 of the color channel value (Krizhevsky et al., 2012). Data augmentation is critical when using ConvNets for image processing and is a common practice for fish classification (Villon et al., 2020, 2021), especially when the available data is limited (Shorten and Khoshgoftaar, 2019). By training the model on variations of the same image, the model is deterred from learning nuanced patterns in the images that can lead to spurious performance, such as the intensity of the background, and encouraged to be robust under variable input conditions. Our preliminary results, as shown in Figure 4, indicate that data augmentation boosted the model’s accuracy by about 2.8%.

### Evaluation Setup for Comparing Classification Performance

In the process of training black-box neural network architectures, it is common to observe higher generalization errors when the amount of training data is small. However, in **HGNN**, we show that by including a biological knowledge-guided loss term (see Equation 1) in the learning objective of neural networks, we can achieve reasonably good generalization performance even in situations where training data are scarce. This is in alignment with the observations made in a previous work by (Jia et al., 2019).

To test for this hypothesis in the context of fish classification, we compared the classification performance of our proposed model to a baseline black-box neural network architecture (termed **Blackbox**-**NN**) comprising of a ResNet of the same size and shape as that of one of the ResNets of our proposed model. Specifically, we compared the performance of **HGNN** and **Blackbox**-**NN** on each of the three data subsets mentioned in Table 1. For each of these subsets, we used 64% of the data for training, 16% for validation, and the remainder for testing. To measure classification performance, we used the f1-score of the correct species class (Tan et al., 2018b). Throughout this paper, we used box plots to show the model’s performance over five random runs of neural network training. To obtain the best-performing neural network models, we performed an explorative Naïve-Bayes approach for hyper-parameter search and fine-tuning. Then, we picked those parameters that performed best on the validation set.

### Tools for Deep Learning Visualization and Assessing Robustness to Adversarial Occlusions

Saliency maps (Simonyan et al., 2014) are heatmaps of the gradients of a neural network model’s output with respect to its input. In other words, a saliency map shows how strongly do changes in pixel values of a certain region of the image cause a change in the species’ probability, highlighting the areas of the image that are most decisive for the classification problem. While other tools, such as GradCAM (Selvaraju et al., 2017), have been used for the same purpose (dos Santos and Gonçalves, 2019), we found saliency maps to be more powerful and capable of detecting the most subtle visual features. Figure 5 shows some examples of saliency maps obtained for **Blackbox**-**NN**. The code we used for generating these saliency maps is inspired by FlashTorch (Ogura and Jain, 2020), an implementation tool based on Guided Back-propagation (Springenberg et al., 2015). As we can see in Figure 5, the baseline model is quite sensitive to different features of the input fish image for different species, including barbels and fins in Figure 5a and the eye in Figure 5b. Saliency maps are also a good debugging tool as they can reveal cases where the model is “cheating” or looking at irrelevant features of the image that are not biologically meaningful for the purpose of fish classification. An example of such a case is presented in Figure 5c, where the model is incorrectly picking up pixels around the note on the label paper in the image as regions with high saliency scores. In this way, saliency maps can be used for “interpreting” the learned features of neural network models.

**Fig. 5:**
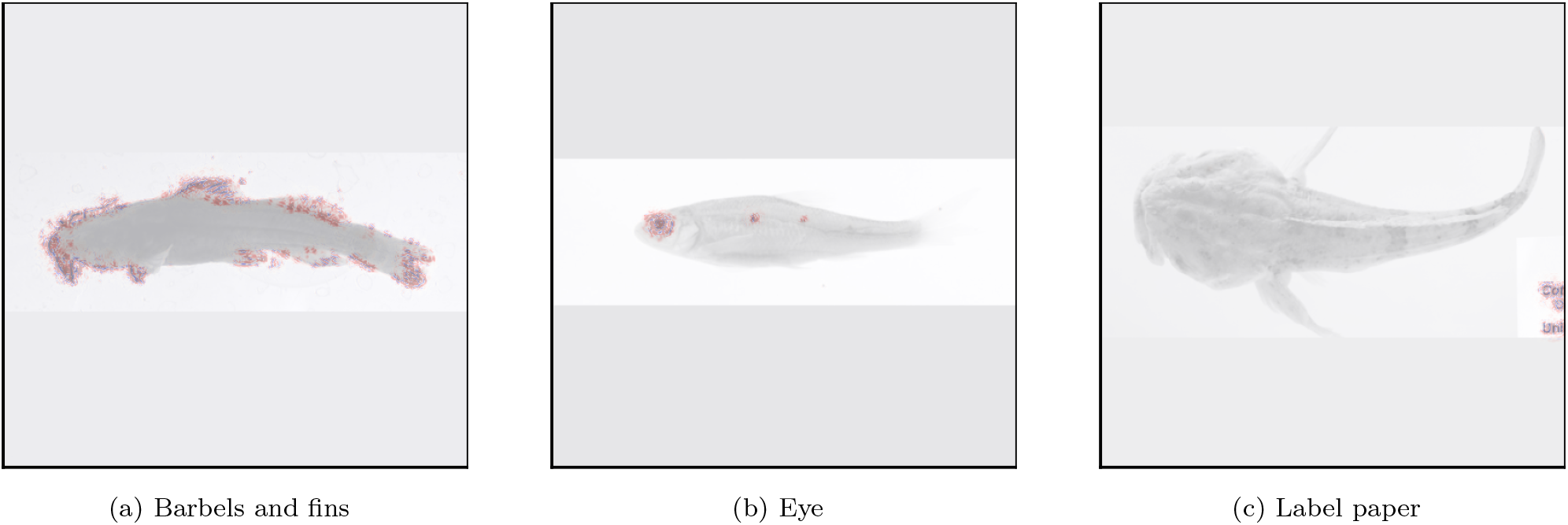
Saliency maps of different fish images obtained for **Blackbox**-**NN**. Pixels in red denote image regions with high saliency scores, indicating higher importance of those regions for fish classification as perceived by the model.

Along with offering interpretability, saliency maps can also be used for investigating the resiliency (or robustness) of neural networks to adversarial occlusions. For example, by occluding regions (or patches) in the input image with high saliency scores, a neural network model’s reliability at making correct predictions can be stress-tested even when it is starved off information from salient image regions. To measure the robustness of a model at every round of adversarial occlusions, we calculated the average probability of the correct class predicted by the model on an input image **x**, averaged over all test images as 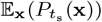. The higher this metric, the less confused the model is about the input. Further, by measuring drops in this metric as a consequence of adversarial occlusions, we can evaluate if a model is too sensitive to selective regions of the input image (with the highest saliency score contributions), which when obstructed can confuse a model into making incorrect predictions. We make use of this metric to assess the robustness of **Blackbox**-**NN** and **HGNN** in our experiments.

## Results

### Effect of Dataset Complexity and Training Size

Figure 6 shows a comparison between **HGNN** and **Blackbox**-**NN** on three subsets of the GLIN dataset: **Easy**/100, **Easy**/50, and **Hard**. Two observations can be made from this figure. First, as datasets become more complex (e.g., the **Hard** dataset) and/or subject to less training data (e.g., the **Easy**/50), the performance of the model deteriorates. Second, and more importantly, the impact of our method is more pronounced exactly when data is scarce and the dataset is complex. As Figure 6 shows, while the median performance of **HGNN** is almost equal to that of **Blackbox**-**NN** for **Easy**/100, which is the easiest of the datasets, the former clearly outperforms the latter on both **Easy**/50 and **Hard**. This highlights our model’s power and ability to compensate for the relative lack of data with respect to dataset complexity by incorporating biological knowledge.

**Fig. 6:**
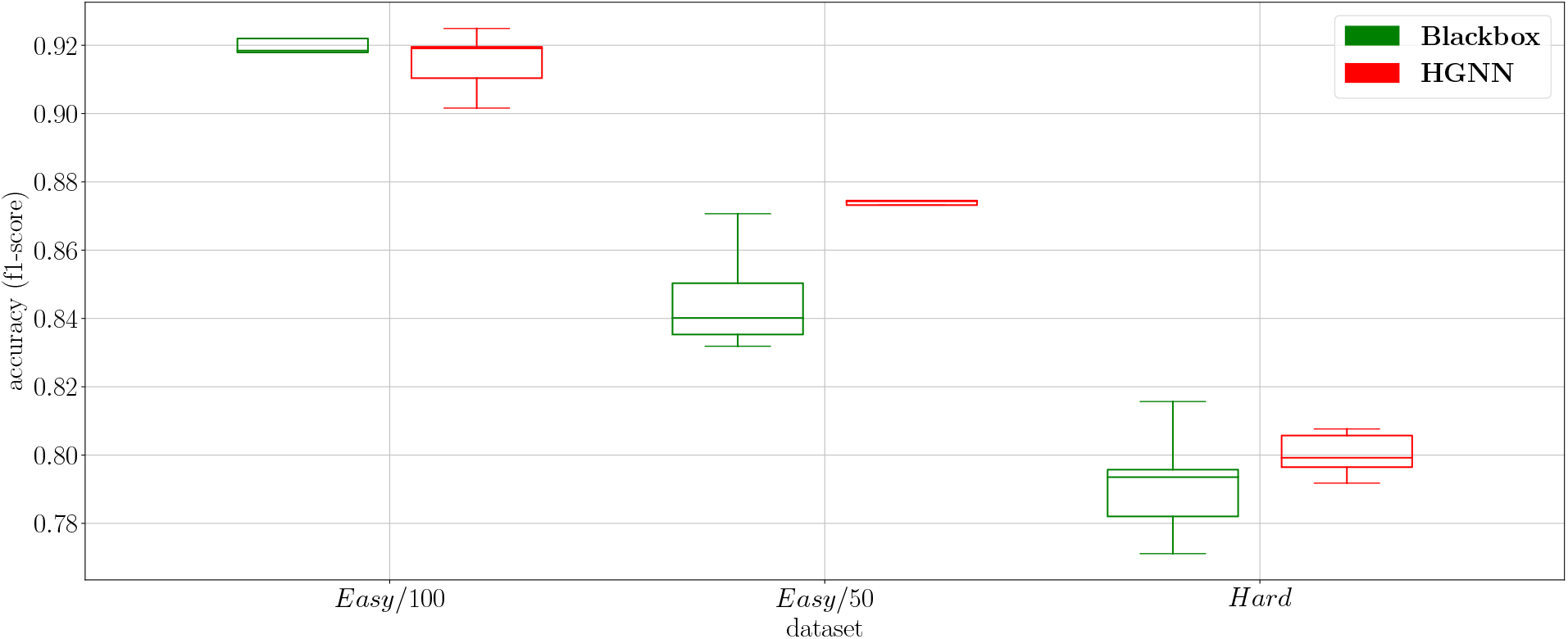
Classification performance across different subsets of the GLIN dataset for **HGNN** and **Blackbox**-**NN**. By definition, as the boxes for the two models do not overlap for **Easy**/50 and **Hard**, it means there is at least 95% confidence (McGill et al., 1978) that the median accuracy of **HGNN** is higher than **Blackbox**-**NN**

### Effect of Adversarial Occlusion

To demonstrate **HGNN**’s resiliency to adversarial occlusions, we iteratively cover regions (or patches) in an image with the highest saliency scores and report the probability of the correct class predicted by the model over the occluded image. Figure 7 shows an example of this process on an illustrative fish specimen from the **Easy**/50 dataset. From left to right, the figure shows a progression from an image with no occlusion towards applying more patches of adversarial occlusions (seen as green square patches) on the same image. Below each image is the model’s predicted probabilities over the 5 most probable species sorted in descending order, for both **HGNN** (top row) and **Blackbox**-**NN** (bottom row). We make a number of observations here. First, all of the saliency maps highlight the features of importance for classifying this fish, namely the eye, nostrils, and the dorsal fin. However, notice that the saliency maps for **HGNN** are slightly different from that of **Blackbox**-**NN**, demonstrating that the two models are not looking at the image in the exact same way (i.e., they have distinct saliency maps). This difference is important for making a fair comparison between the two models. Second, even when there is no occlusion, while **Blackbox**-**NN** makes the correct prediction, its probability of the correct species class is significantly lower than that of **HGNN**’s. This demonstrates **HGNN**’s ability to extract more useful and generalizable features from images for fish classification. Third, after applying two patches of occlusions (in the middle column), we notice that even though both models get the species right, **Blackbox**-**NN**’s second guess is not within the correct genus. Finally, and most importantly, after applying four patches of occlusions (in rightmost column), we notice that while both models start predicting the wrong class, **HGNN** is still within the correct genus, while **Blackbox**-**NN** is not. It follows that the **Blackbox**-**NN** model is not learning phylogenetic features that could be used in other tasks, such as trait segmentation. To drive this point home, we automate this process for the entire dataset and compute the average predicted probability of the correct class across all images, as a function of the number of adversarial occlusions applied to the images. Table 2 reports the results for each number of patches ranging from 0 (no occlusion) to 4. We can see that **HGNN** shows higher average probability of the correct class across all number of patches in comparison with **Blackbox**-**NN**. This demonstrates **HGNN**’s ability to generalize and handle image imperfections better, especially when the most informative (or salient) regions of the image are occluded.

**Fig. 7:**
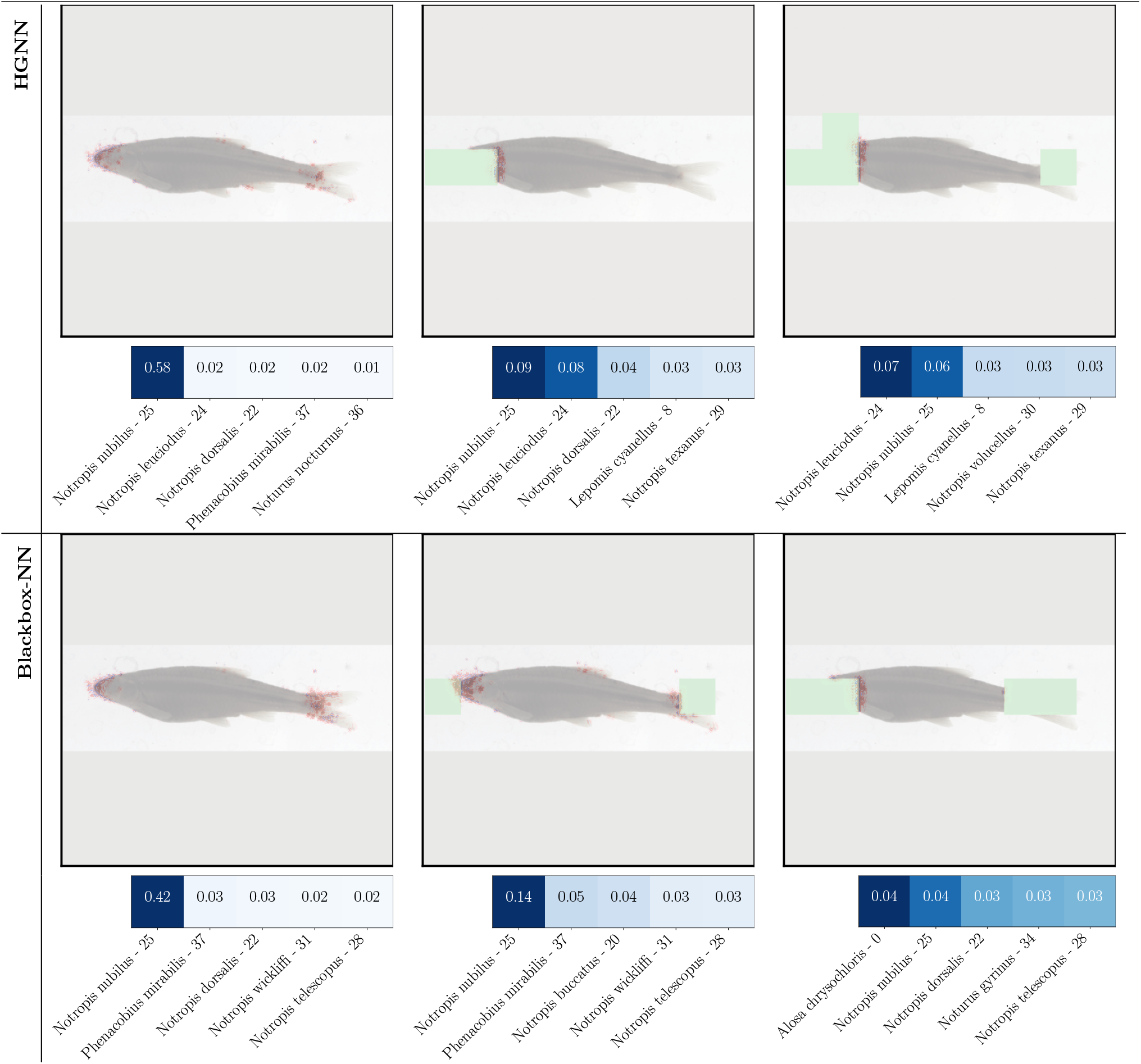
Saliency maps showing the effect of adversarial occlusions (shown as green square patches) on the predicted probabilities of the species class produced by **HGNN** (top row) and **Blackbox**-**NN** (bottom row) on an example fish image. The left-most column corresponds to the case with no occlusion, while the number of occlusions increase as we go from left column to the middle column (2 patches) to the right-most column (4 patches).

**Table 2.**
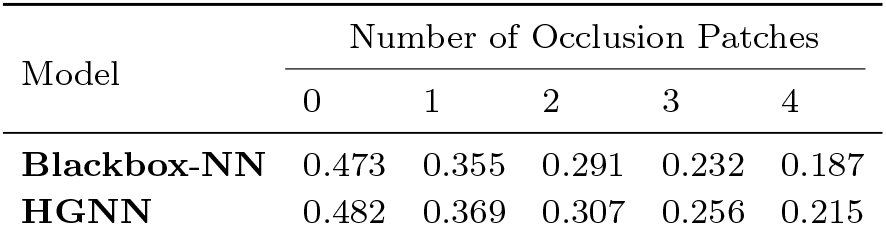
Average probability of the correct species class predicted by **Blackbox**-**NN** and **HGNN** over **Easy**/50, as a function of the number of adversarial occlusions applied to every image. From left to right, we start with non-occluded images and progressively add more patches of occlusions.

| Model | Number of Occlusion Patches |  |  |  |  |
| --- | --- | --- | --- | --- | --- |
|  | 0 | 1 | 2 | 3 | 4 |
| **Blackbox-NN** | 0.473 | 0.355 | 0.291 | 0.232 | 0.187 |
| **HGNN** | 0.482 | 0.369 | 0.307 | 0.256 | 0.215 |

## Discussion

In this paper, we have shown that embedding the hierarchical taxonomy of the genus and species classes in the design and learning of neural networks leads to solutions with better generalization, superior accuracy, and better resiliency to adversarial occlusions. Most of the deep learning methods currently in the literature perform tasks without learning biologically-relevant features. Our proposed method leverages a particularly important aspect of species classification—the hierarchical arrangement of taxon names—which improves model interpretability and biological-validity. The aim of our method is to provide biologists not only with the correct classification, but also with a plausible one when it fails.

An ultimate goal of this research is to augment biological information on the connections among phenotype, genotype and environment into deep learning, so that an understanding of genealogical relationships among species is discovered by our neural networks. While we have not fully investigated these relationships here, a future direction of our project is to explore how the anatomical features of species learned by our models relate to the environments the species were collected from and how closely related the species are. This would increase understanding of how the environment and genealogy shape the phenotypes of species. Moreover, we plan to investigate how such learned features aid us in other relevant tasks, such as segmenting the phenotypic traits of species. Finally, we also plan to exploit other forms of hierarchical information such as phylogenetic tree-based distances among species to better understand how this informs biologically-informed neural network feature learning.

Recent advances in image computation are enabling automated methods of extracting phenotypic data from specimen images. We hope that our present framework for leveraging biological information in training machine learning models will have a direct impact on several biologically relevant computer vision tasks, including species detection (Li et al., 2016), tracking and counting (Spampinato et al., 2008), segmentation (Chuang et al., 2013; Yao et al., 2013), and classification (Ding et al., 2017; Rathi et al., 2018; Sarigul, 2017). This automation effort is essential as manual annotation is laborious and requires expertise (Villon et al., 2020), especially with the large amount of data that has become recently available (Ditria et al., 2020a). Moreover, it has been shown that automation can be more accurate than human annotation (Ditria et al., 2020b).

In this paper, we have focused on teleost fishes as a model system for species classification due to their high diversity and importance economically and scientifically. Fishes are the targets of recreation (Arlinghaus and Cooke, 2009), aquaculture and fisheries (Lynch et al., 2016), and conservation (Arthington et al., 2016). Fishes make up more than half of all vertebrates and they play critical roles in Earth ecosystems (Near et al., 2012; Villon et al., 2020). However, our framework of **HGNN** is quite generic and can be potentially applied to incorporate hierarchical knowledge into machine learning models for a broad variety of other biological problems involving phenotypic trait discovery and understanding in other taxonomic groups.

## Acknowledgment

This work is supported by the National Science Foundation through Harnessing the Data Revolution Ideas Lab program awards 1940322 (Bart); 1940233 (Greenberg); 2022042 (Mabee); 1940247 (Karpatne) and 1939505 (Maga), and also a startup allocation from the XSEDE (TG-DEB200005).

## Data Availability

For this work, and in an effort to create a diverse and statistically substantial dataset, we aggregated more than 60,000 images of fish specimens from five ichthyological research collections (Field Museum of Natural History http://www.tubri.org/HDR/FMNH/, Illinois Natural History Survey http://www.tubri.org/HDR/INHS/, J. F. Bell Museum of Natural History (http://www.tubri.org/HDR/JFBM/), Ohio State University Museum of Biological Diversity (http://www.tubri.org/HDR/OSUM/), and the University of Wisconsin-Madison Zoological Museum (http://www.tubri.org/HDR/UWZM/)) that participated in the Great Lakes Invasives Network Project (GLIN) ^1^. The GLIN project is digitizing 1.73 million historical biological specimens representing 2,550 species, including fishes, clams, snails, mussels, algae and plants that are potentially invasive to the Great Lakes Region of the U.S. The computer code used for running our experiments is found at (Elhamod, 2021).

## Authors’ contribution

M.E designed the methodology from machine learning perspective and conducted and analysed the experiments. K.D, A.M.M and B.A pre-processed the data and helped design the experiments from phylogeny perspective. Y.B collected and labelled the data. H.B, P.M and W.D critiqued the methodology and provided suggestions to improve it. They also helped in selecting the set of species to work on and the general direction of research to explore. J.L and J.G helped setting up the pipeline for managing image metadata and creating workflows for model deployment. A.K provided overall supervision across all tasks conducted in this work. All authors contributed to writing the manuscript and gave final approval for publication.

## Footnotes

1 http://greatlakesinvasives.org/

