## Supplemental File for "Hierarchy-guided Neural Networks for Species Classification" (from another version of this preprint: version 2, 2021-10-05)

[1]Virginia Tech, Blacksburg, 24060, VA, USA, [2]Seattle Children's Research Institute, Seattle, 98199, WA, USA, [3]University of Washington, Seattle, 98115, WA, USA, [4]Tulane University, New Orleans, 70118, LA, USA, [5]National Ecological Observatory Network, Battelle, Boulder, 80304, CO, USA, [6]University of California, Irvine, Irvine, 92623, CA, USA, [7]Drexel University, Philadelphia, 1910, PA, USA and [8]University of Virginia, Charlottesville, 22904, VA, USA

### Abstract

1. Species classification is an important task that is the foundation of industrial, commercial, ecological, and scientific applications involving the study of species distributions, dynamics, and evolution.
2. While conventional approaches for this task use off-the-shelf machine learning (ML) methods such as existing Convolutional Neural Network (ConvNet) architectures, there is an opportunity to inform the ConvNet architecture using our knowledge of biological hierarchies among taxonomic classes.
3. In this work, we propose a new approach for species classification termed Hierarchy-Guided Neural Network (**HGNN**), which infuses hierarchical taxonomic information into the neural network's training to guide the structure and relationships among the extracted features. We perform extensive experiments on an illustrative use-case of classifying fish species to demonstrate that **HGNN** outperforms conventional ConvNet models in terms of classification accuracy, especially under scarce training data conditions.
4. We also observe that **HGNN** shows better resilience to adversarial occlusions, when some of the most informative patch regions of the image are intentionally blocked and their effect on classification accuracy is studied.

### Assessments of Monophyly

**Table S1.** List of the 24 genera from which species were examined in this study. Most of the genera (21) are monophyletic, and thus their taxonomy mirrors their evolutionary relationships. Three of the 24 genera, however, are not monophyletic (i.e., some of the species in the genus are more closely related to species from other genera, than they are to other species in the genus). Here the taxonomic hierarchy does not capture similarity due to common ancestry.

| Genus | Reference supporting monophyly | Monophyly |
|---|---|---|
| *Alosa* | (Bowen et al., 2008) | Monophyletic |
| *Carassius* | (Rylková et al., 2010) | Monophyletic |
| *Chrosomus* | (Schönhuth et al., 2018) | Monophyletic |
| *Cottus* | (Kinziger et al., 2005) | Included species are members of monophyletic *Uranidea Clade* |
| *Cyprinella* | (Schönhuth et al., 2018) | Monophyletic |
| *Cyprinus* | (Chen et al., 2017) | Monophyletic, tetraploidy |
| *Enneacanthus* | (Near et al., 2004) | Monophyletic |
| *Esox* | (Grande et al., 2004) | Monophyletic |
| *Fundulus* | (Rodgers, 2017) | Not Monophyletic |
| *Gambusia* | (Hrbek et al., 2007) | Monophyletic |
| *Hybopsis* | (Schönhuth et al., 2018) | Monophyletic |
| *Lepisosteus* | (Normark et al., 1991) | Monophyletic |
| *Lepomis* | (Harris et al., 2005) | Monophyletic |
| *Luxilus* (a.k.a *Luxilis*) | (Schönhuth et al., 2018) | Not monophyletic |
| *Lythrurus* | (Schönhuth et al., 2018) | Monophyletic |
| *Morone* | (Waldman, 1987) | Monophyletic |
| *Neogobius* | (Neilson and Stepien, 2009) | Monophyletic |
| *Notropis* | (Schönhuth et al., 2018) | Not monophyletic |
| *Noturus* | (Lundberg, 1970) and (Grady, 1993) | Monophyletic |
| *Oncorhynchus* | (Campbell et al., 2020) | Monophyletic |
| *Opsopoeodus* | (Schönhuth et al., 2018) | Monophyletic with *Pimephales* |
| *Osmerus* | (Taylor and Dodson, 1994) | Monophyletic |
| *Perca* | (Sloss et al., 2004) | Monophyletic |
| *Phenacobius* | (Schönhuth et al., 2018) | Monophyletic |
| *Salmo* | (Campbell et al., 2020) | Monophyletic |

### Species Statistics

**Table S2.** List of all the species in the **Easy**/50 dataset, with the number of samples per species.

| Species | Count |
|---|---|
| *Alosa chrysochloris* | 50 |
| *Carassius auratus* | 50 |
| *Cyprinus carpio* | 50 |
| *Esox americanus* | 50 |
| *Gambusia affinis* | 50 |
| *Lepisosteus osseus* | 50 |
| *Lepisosteus platostomus* | 50 |
| *Lepomis auritus* | 50 |
| *Lepomis cyanellus* | 50 |
| *Lepomis gibbosus* | 50 |
| *Lepomis gulosus* | 50 |
| *Lepomis humilis* | 50 |
| *Lepomis macrochirus* | 50 |
| *Lepomis megalotis* | 50 |
| *Lepomis microlophus* | 50 |
| *Morone chrysops* | 50 |
| *Morone mississippiensis* | 50 |
| *Notropis atherinoides* | 50 |
| *Notropis blennius* | 50 |
| *Notropis boops* | 50 |
| *Notropis buccatus* | 50 |
| *Notropis buchanani* | 50 |
| *Notropis dorsalis* | 50 |
| *Notropis hudsonius* | 50 |
| *Notropis leuciodus* | 50 |
| *Notropis nubilus* | 50 |
| *Notropis percobromus* | 50 |
| *Notropis stramineus* | 50 |
| *Notropis telescopus* | 50 |
| *Notropis texanus* | 50 |
| *Notropis volucellus* | 50 |
| *Notropis wickliffi* | 50 |
| *Noturus exilis* | 50 |
| *Noturus flavus* | 50 |
| *Noturus gyrinus* | 50 |
| *Noturus miurus* | 50 |
| *Noturus nocturnus* | 50 |
| *Phenacobius mirabilis* | 50 |

**Table S3.** List of all the species in the **Easy**/100 dataset, with the number of samples per species.

| Species | Count |
|---|---|
| *Alosa chrysochloris* | 63 |
| *Carassius auratus* | 99 |
| *Cyprinus carpio* | 100 |
| *Esox americanus* | 100 |
| *Gambusia affinis* | 100 |
| *Lepisosteus osseus* | 100 |
| *Lepisosteus platostomus* | 100 |
| *Lepomis auritus* | 100 |
| *Lepomis cyanellus* | 100 |
| *Lepomis gibbosus* | 100 |
| *Lepomis gulosus* | 100 |
| *Lepomis humilis* | 100 |
| *Lepomis macrochirus* | 100 |
| *Lepomis megalotis* | 100 |
| *Lepomis microlophus* | 100 |
| *Morone chrysops* | 100 |
| *Morone mississippiensis* | 100 |
| *Notropis atherinoides* | 100 |
| *Notropis blennius* | 100 |
| *Notropis boops* | 100 |
| *Notropis buccatus* | 100 |
| *Notropis buchanani* | 100 |
| *Notropis dorsalis* | 100 |
| *Notropis hudsonius* | 100 |
| *Notropis leuciodus* | 100 |
| *Notropis nubilus* | 100 |
| *Notropis percobromus* | 100 |
| *Notropis stramineus* | 100 |
| *Notropis telescopus* | 100 |
| *Notropis texanus* | 100 |
| *Notropis volucellus* | 100 |
| *Notropis wickliffi* | 100 |
| *Noturus exilis* | 100 |
| *Noturus flavus* | 100 |
| *Noturus gyrinus* | 100 |
| *Noturus miurus* | 100 |
| *Noturus nocturnus* | 100 |
| *Phenacobius mirabilis* | 100 |

**Table S4.** List of all the species in the **Hard** dataset, with the number of samples per species.

| Species | Count | Species | Count |
|---|---|---|---|
| *Alosa chrysochloris* | 50 | *Notropis anogenus* | 50 |
| *Alosa pseudoharengus* | 50 | *Notropis ariommus* | 33 |
| *Carassius auratus* | 50 | *Notropis atherinoides* | 50 |
| *Chrosomus eos* | 50 | *Notropis bifrenatus* | 50 |
| *Chrosomus erythrogaster* | 50 | *Notropis blennius* | 50 |
| *Chrosomus neogaeus* | 40 | *Notropis boops* | 50 |
| *Cottus bairdii* | 50 | *Notropis buccatus* | 50 |
| *Cottus carolinae* | 50 | *Notropis buchanani* | 50 |
| *Cottus cognatus* | 50 | *Notropis chalybaeus* | 50 |
| *Cottus perplexus* | 50 | *Notropis chihuahua* | 50 |
| *Cottus rhotheus* | 33 | *Notropis cummingsae* | 33 |
| *Cyprinella camura* | 50 | *Notropis dorsalis* | 50 |
| *Cyprinella galactura* | 50 | *Notropis edwardraneyi* | 50 |
| *Cyprinella lutrensis* | 50 | *Notropis greenei* | 50 |
| *Cyprinella spiloptera* | 50 | *Notropis heterodon* | 50 |
| *Cyprinella venusta* | 31 | *Notropis heterolepis* | 50 |
| *Cyprinella whipplei* | 50 | *Notropis hudsonius* | 50 |
| *Cyprinus carpio* | 50 | *Notropis leuciodus* | 50 |
| *Enneacanthus gloriosus* | 50 | *Notropis longirostris* | 46 |
| *Esox lucius* | 50 | *Notropis lutipinnis* | 49 |
| *Esox masquinongy* | 46 | *Notropis micropteryx* | 50 |
| *Esox niger* | 44 | *Notropis nazas* | 50 |
| *Fundulus catenatus* | 50 | *Notropis nubilus* | 50 |
| *Gambusia alvarezi* | 34 | *Notropis percobromus* | 50 |
| *Gambusia bonita* | 50 | *Notropis photogenis* | 50 |
| *Gambusia parismina* | 40 | *Notropis procne* | 50 |
| *Hybopsis amblops* | 50 | *Notropis rubellus* | 50 |
| *Lepisosteus oculatus* | 50 | *Notropis rubricroceus* | 38 |
| *Lepisosteus osseus* | 50 | *Notropis spectrunculus* | 34 |
| *Lepisosteus platostomus* | 45 | *Notropis stilbius* | 39 |
| *Lepomis auritus* | 50 | *Notropis stramineus* | 50 |
| *Lepomis cyanellus* | 50 | *Notropis telescopus* | 50 |
| *Lepomis gibbosus* | 50 | *Notropis texanus* | 50 |
| *Lepomis gulosus* | 50 | *Notropis topeka* | 50 |
| *Lepomis humilis* | 50 | *Notropis volucellus* | 50 |
| *Lepomis macrochirus* | 50 | *Notropis wickliffi* | 50 |
| *Lepomis marginatus* | 50 | *Notropis xaenocephalus* | 42 |
| *Lepomis megalotis* | 50 | *Noturus albater* | 50 |
| *Lepomis microlophus* | 50 | *Noturus eleutherus* | 50 |
| *Lepomis miniatus* | 50 | *Noturus exilis* | 44 |
| *Lepomis peltastes* | 50 | *Noturus hildebrandi* | 41 |
| *Lepomis punctatus* | 50 | *Noturus insignis* | 50 |
| *Lepomis symmetricus* | 34 | *Noturus leptacanthus* | 50 |
| *Luxilua zonatus* | 49 | *Oncorhynchus kisutch* | 48 |
| *Luxilus chrysocephalus* | 50 | *Oncorhynchus mykiss* | 50 |
| *Luxilus coccogenis* | 50 | *Oncorhynchus tshawytscha* | 50 |
| *Luxilus cornutus* | 50 | *Opsopoeodus emiliae* | 39 |
| *Lythrurus umbratilis* | 50 | *Osmerus mordax* | 50 |
| *Lythrutus ardens* | 50 | *Perca flavescens* | 50 |
| *Morone americana* | 50 | *Phenacobius uranops* | 50 |
| *Neogobius melanostomus* | 50 | *Salmo trutta* | 50 |

Rachel Rodgers. *Phylogenomic Analysis of Fundulidae Using RNA-Sequencing Data.* PhD thesis, Southern Illinois University at Edwardsville, 2017.

Kateřina Rylková, Lukáš Kalous, Vendula Šlechtová, and Jörg Bohlen. Many branches, one root: First evidence for a monophyly of the morphologically highly diverse goldfish (carassius auratus). *Aquaculture*, 302(1):36 – 41, 2010. ISSN 0044-8486. doi: https://doi.org/10.1016/j.aquaculture.2010.02.003. URL http://www.sciencedirect.com/science/article/pii/S0044848610000918.

Susana Schönhuth, Jasna Vukić, Radek Šanda, Lei Yang, and Richard L. Mayden. Phylogenetic relationships and classification of the holarctic family leuciscidae (cypriniformes: Cyprinoidei). *Molecular Phylogenetics and Evolution*, 127:781 – 799, 2018. ISSN 1055-7903. doi: https://doi.org/10.1016/j.ympev.2018.06.026. URL http://www.sciencedirect.com/science/article/pii/S1055790317308618.

Brian L Sloss, Neil Billington, and Brooks M Burr. A molecular phylogeny of the percidae (teleostei, perciformes) based on mitochondrial dna sequence. *Molecular Phylogenetics and Evolution*, 32(2):545 – 562, 2004. ISSN 1055-7903. doi: https://doi.org/10.1016/j.ympev.2004.01.011. URL http://www.sciencedirect.com/science/article/pii/S105579030400048X.

E. B. Taylor and J. J. Dodson. A molecular analysis of relationships and biogeography within a species complex of holarctic fish (genus osmerus). *Molecular Ecology*, 3(3):235–248, 1994. doi: https://doi.org/10.1111/j.1365-294X.1994.tb00057.x. URL https://onlinelibrary.wiley.com/doi/abs/10.1111/j.1365-294X.1994.tb00057.x.

JR Waldman. Systematics of morone (pisces: Moronidae), with notes on the lower percoids (distribution, cladistics, striped bass, siniperca, centropomidae). 1987.
